# Molecular structure discovery for untargeted metabolomics using biotransformation rules and global molecular networking

**DOI:** 10.1101/2024.02.04.578795

**Authors:** Margaret R. Martin, Wout Bittremieux, Soha Hassoun

## Abstract

Although untargeted mass spectrometry-based metabolomics is crucial for understanding life’s molecular underpinnings, its effectiveness is hampered by low annotation rates of the generated tandem mass spectra. To address this issue, we introduce a novel data-driven approach, Biotransformation-based Annotation Method (BAM), that leverages molecular structural similarities inherent in biochemical reactions. BAM operates by applying biotransformation rules to known ‘anchor’ molecules, which exhibit high spectral similarity to unknown spectra, thereby hypothesizing and ranking potential structures for the corresponding ‘suspect’ molecule. BAM’s effectiveness is demonstrated by its success in annotating suspect spectra in a global molecular network comprising hundreds of millions of spectra. BAM was able to assign correct molecular structures to 24.2 % of examined anchor-suspect cases, thereby demonstrating remarkable advancement in metabolite annotation.

---

Metabolomics, empowered by untargeted mass spectrometry (MS), is pivotal in advancing phenotyping and biomarker discovery by profiling thousands of small molecules using tandem mass spectrometry (MS/MS). However, a critical challenge hampers its full potential: the low MS/MS spectrum annotation rate. Despite technological advancements and expanding spectral libraries, a significant portion of spectra remain unidentified. For example, only a small minority of spectra in key public data repositories like GNPS/MassIVE [1] has been successfully annotated [2].

This gap in annotation is particularly pressing given the crucial role of metabolites as direct products of metabolic processes. Some methods attempt to address this by leveraging the biochemical transformations and structural similarities intrinsic to metabolites [3, 4]. These approaches utilize MS/MS spectra and empirical mass differences to predict structural relationships and integrate reactants from metabolic networks to improve annotation confidence [5]. However, despite these efforts, existing methodologies fall short in effectively interpreting the vast majority of MS/MS data produced in untargeted metabolomics experiments. This limitation not only restricts biological insights that could be gleaned from expensive and critical studies, but also hinders a deeper understanding of metabolism and its products. Thus, there is an urgent need for innovative approaches that can bridge this gap in MS/MS spectrum annotation, thus enabling more comprehensive exploration and interpretation of the metabolome in untargeted metabolomics studies.

To address this problem, we have developed the Biotransformation-based Annotation Method (BAM). At its core, BAM leverages the principle that substrates and products of biochemical reactions often share molecular substructures, leading to high spectral similarity [6]. This spectral similarity, when analyzed within a biological context, provides a more biologically relevant and comprehensive approach to annotation compared to traditional methods that rely solely on spectral similarity. BAM operates by identifying and applying biotransformation rules to an ‘anchor’ molecule—a known compound whose MS/MS spectrum exhibits high similarity to a given query spectrum. These rules, drawn from extensive biochemical databases like RetroRules [7] and KEGG [8], encapsulate the promiscuous behavior of enzymes towards substrates. Specifically, they include a molecular substructure specifying a reaction center and the structural changes occurring therein. BAM systematically employs these rules to generate and rank potential molecular structures (‘derivatives’) for the ‘suspect’—the unknown compound associated with the query spectrum. To do so, BAM makes use of PROXIMAL2 [9] to apply the biotransformation rules to the anchor molecule, identifying suitable reaction centers for the proposed structural changes. Next, the graph neural network-based GNN-SOM tool is used to predict the site-of-metabolism and rank these putative derivatives based on the likelihood of each atom being a reaction center [10].

An example application of BAM on three queried suspects is demonstrated (figure 1). All suspect spectra exhibit high spectral similarity to the same anchor, cholic acid (figure 1a). BAM identifies relevant mass differences between the suspects and the anchor, selects appropriate biotransformation rules corresponding to these mass differences (figure 1b), and applies them to generate and rank multiple product candidates (figure 1c). In this case, BAM was able to successfully identify the true structures of the suspects, showcasing its potential in discovering new molecules corresponding to previously unknown MS/MS spectra. Consequently, BAM not only enhances the accuracy of compound identification from untargeted metabolomics data, but it also enables the discovery of novel molecules, thereby enriching our comprehension of metabolic processes and their vast array of products.

**Figure 1:**
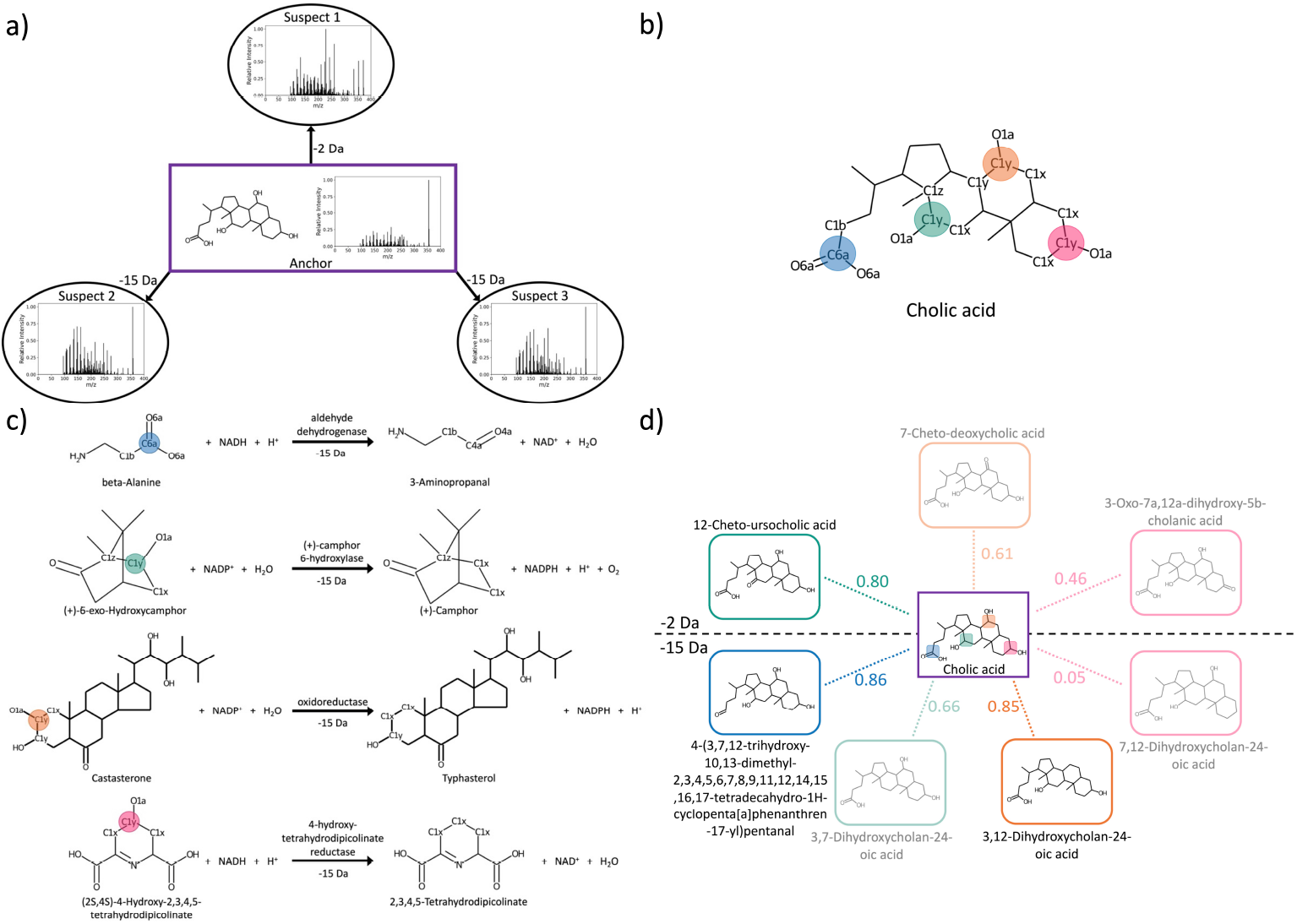
Demonstration of BAM using cholic acid as an anchor molecule to annotate three suspect spectra. **(a)** The anchor molecule, cholic acid, has high spectral similarity with three unidentified suspect spectra. The precursor *m*/*z* differences between the anchor and suspects correspond to −15 Da and −2 Da. **(b)** Biotransformation rules are selected by the observed mass differences and are only applied if the corresponding reaction center is exhibited in the anchor structure. Selected biotransformations rules are applied to four different reaction centers, which are highlighted by different colors here. The KEGG atom type of the reaction centers as well as the neighboring atoms are displayed as well. **(c)** The four relevant biotransformation rules that exhibit a mass difference of −15 Da are identified. Each biotransformation rule may be applied to the corresponding reaction center of cholic acid, as indicated by color. **(d)** Application of BAM: The four depicted biotransformation rules are applied to cholic acid to generate four derivative candidates for suspect 2 and 3 that have a mass difference of −15 Da. Three other biotransformation rules are applied to cholic acid to generate three derivative candidates for suspect 1 that have a mass difference of −2 Da. The generated derivative candidates are ranked based on the predicted likelihood of each atom within the anchor molecule serving as a reaction center. The figure distinguishes candidates derived from the −2 Da rules (above the dashed line) and those from the −15 Da rules (below the dashed line). The likelihood of each product structure being the correct annotation is indicated along corresponding colored dotted lines. The true structures of the three suspects are the top-ranked candidates (displayed without shading). The top-ranked candidate for the −2 Da mass difference— 12-Cheto-ursocholic acid—matches the actual structure of suspect 1. Similarly, the two highest-ranked candidates for the −15 Da mass difference—4-(3,7,12-trihydroxy-10,13-dimethyl-2,3,4,5,6,7,8,9,11,12,14,15,16,17-tetradecahydro-1H-cyclopenta[a]phenanthren-17-yl)pentanal and 3,12-Dihydroxycholan-24-oic acid—both displaying significant structural similarity, match the true structures of the remaining two suspects.

Next, we applied BAM to a global molecular network that was recently compiled from hundreds of millions of public MS/MS spectra [11] available on GNPS/MassIVE [1], MetaboLights [12], and Metabolomics Workbench [13]. From this global molecular network, a set of 30 184 structurally unique, annotated spectrum pairs was derived, yielding 60 368 potential anchor–suspect pairs. Additionally, we utilized RetroRules as the basis for extracting biotransformation rules [7]—yielding 42 307 unique rules—covering a wide range of possible biochemical transformations. Here, 41 % of the rules overlapped with the mass differences exhibited by the anchor–suspect pairs, while 98 % of mass differences are covered by the biotransformation rules.

Applying BAM resulted in 17 271 anchors generating one or more derivatives. Among these, BAM successfully generated the correct suspect structure in 4171 cases, achieving a recall rate of 24.2 % when a derivative is generated. Notably, when the structural similarity between the anchor and suspect was restricted to above 0.9 Tanimoto similarity, the recall rate increased significantly to 37.7 %, highlighting BAM’s effectiveness for similar molecules (figure 2a). Furthermore, BAM demonstrated impressive accuracy in ranking the correct derivatives, achieving an average rank of 2.1 and ranking the correct suspect first in 57.7 % of cases (figure 2b).

**Figure 2:**
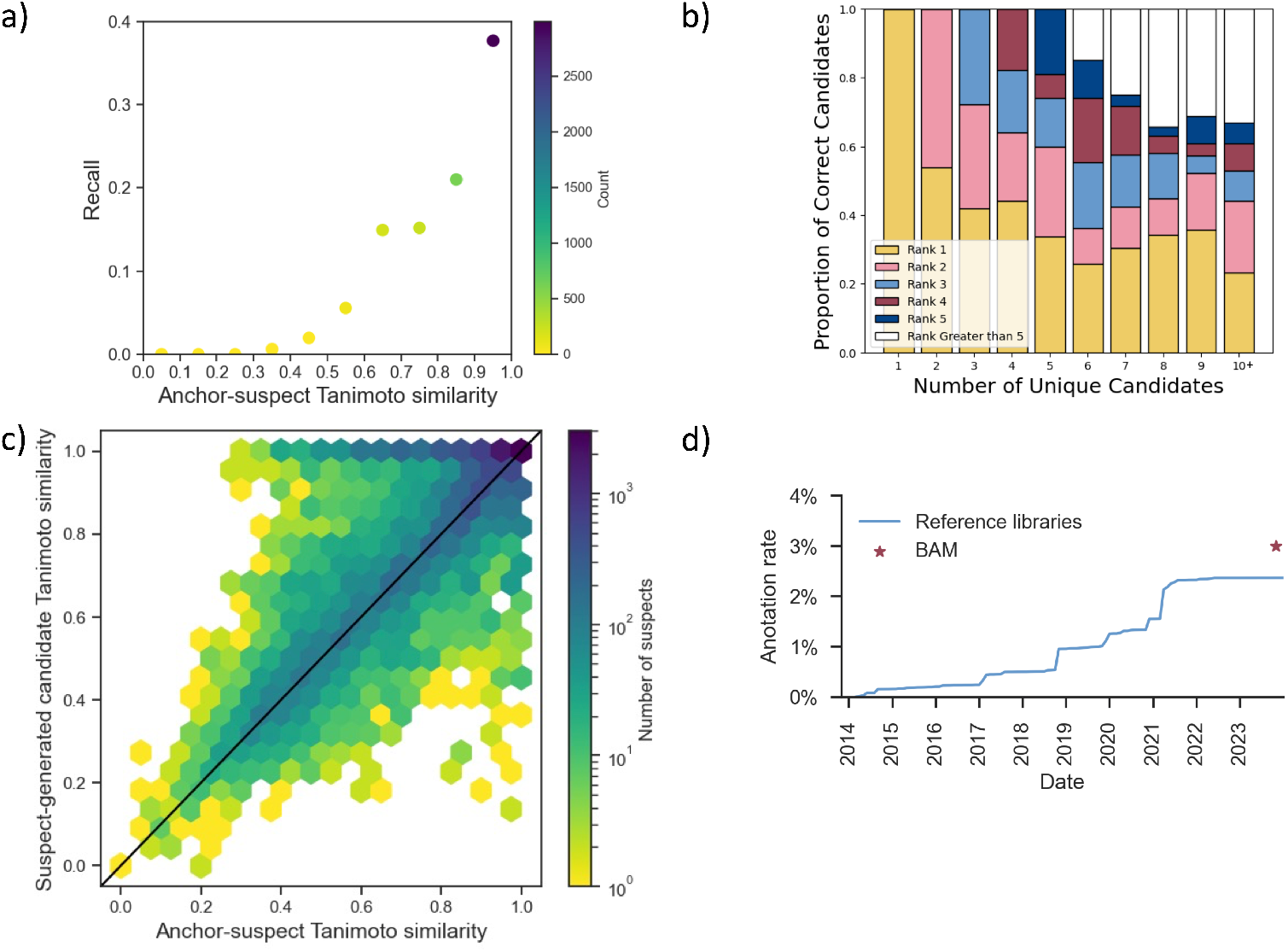
Performance evaluation of BAM. **(a)** Recall rate of BAM in function of the structural similarity between the anchor and suspect molecules. The recall and the count capture the proportion and the number of queries for which BAM generated the true structure when a derivative is generated, respectively. The count is represented by the color of each point. **(b)** Ranking of the candidates from anchor spectra for which derivatives could be generated. **(c)** Anchor–suspect Tanimoto similarity versus suspect–generated candidate Tanimoto similarity. In 74.0 % of cases (observations on or above the diagonal), BAM generated a candidate more similar to the suspect than the anchor. The BAM candidates therefore mark an improvement in structural hypotheses for metabolite annotation of previously unknown MS/MS spectra. **(d)** Contribution offered by BAM in the context of the historical increase in MS/MS spectrum annotation rates [2], as determined by spectral library searching performed as part of the systematic living data reanalysis on GNPS [1].

An analysis of the derivatives generated by BAM revealed that in 74.0 % of cases, the candidates were more similar or equally similar to the suspect than to the anchor molecule (figure 2c). This indicates that BAM can not only identify potential suspect structures, but also improves upon structural hypotheses for metabolite annotation than standard molecular networking or analog searching. Such a high degree of similarity enhancement is a testament to BAM’s ability to accurately interpret and utilize biochemical transformations for MS/MS spectrum annotation.

When considering the increase in spectrum annotation rate afforded by BAM, its potential becomes even more apparent. As each node in the global molecular network represents a number of experimental spectra, the 4171 correctly annotated queries map to 219 985 spectra across 466 GNPS/MassIVE datasets. The BAM approach is therefore widely applicable and can potentially provide novel insights for a diverse range of biological studies, underscoring its broad relevance and effectiveness. Extrapolating the recall rate from these findings to all anchor–suspect pairs in the global molecular network, BAM could annotate approximately 1 135 000 previously unknown MS/MS spectra from 1129 datasets in GNPS/MassIVE, which increases the overall annotation rate from 2.3% to 3% (figure 2d). This substantial enhancement in annotation capabilities marks a significant leap forward in metabolite discovery and understanding, as BAM not only boosts the recall rate for correct structural assignments but also offers richer structural hypotheses for previously unannotated metabolites.

Looking ahead, there are opportunities to further refine and enhance BAM’s performance. One promising avenue is the extraction of additional biotransformation rules from specific biochemical reactions, particularly those most relevant to the biological samples under study. Improvements in the ranking accuracy of BAM could also be achieved by finetuning the GNN-SOM model using data from RetroRules. Furthermore, the integration of MS/MS data during scoring, along with the consideration of multiple anchor spectra, could improve the generation and ranking of candidate structures. For instance, candidates for a suspect derived from different anchors could be ranked based on a consensus metric, providing a more robust and accurate annotation.

In conclusion, BAM’s application to a global molecular network signals an important step forward in metabolomics, enabling discovery and annotation of novel analog molecules in a data-driven fashion, rather than being restricted by exact matching to existing spectral libraries with limited coverage. Its methodology, which harnesses vast public MS/MS data and biotransformation rules, paves the way for deeper insights into metabolic processes and significantly enriches the field of metabolite annotation. With ongoing enhancements and refinements, BAM is poised to unlock even more potential in the exploration and understanding of the complex world of metabolomics.

## Online Methods

BAM consist of three steps:

1. For a given query spectrum, determine an annotated anchor spectrum based on high spectral similarity.
2. Apply relevant biotransformation rules that match the observed mass difference between the anchor and suspect to generate candidate structures—derivatives—for the query spectrum.
3. Rank the derivatives based on the likelihoods of the applied biotransformation.

Next, we will describe each of these steps in detail.

### Identification of anchor molecules based on spectral similarity

In the initial step, for a given query MS/MS spectrum, BAM identifies an annotated anchor spectrum that closely resembles the query spectrum in terms of spectral similarity. This can be achieved through various methods, including analog spectral library searching, molecular networking [14], or by querying against public data repositories such as GNPS/MassIVE [1] (e.g. using MASST [15]) or MoNA [16]. Here, we utilized a global molecular network that was recently constructed from 521 million MS/MS spectra in 1335 public projects deposited to the GNPS/MassIVE [1], MetaboLights [12], and Metabolomics Workbench [13] public data repositories. This network comprises 8 543 020 nodes, including 454 091 spectra that could be annotated using standard spectral library searching against the default GNPS spectral libraries, forming a large-scale molecular network of 13 179 147 spectrum pair edges [11].

From this, we identified paired annotated compounds with high spectral similarity (modified cosine similarity above 0.8). While the majority of the edges in the molecular network (8 599 249) connect pairs of unannotated nodes, only a small number of edges (616 602) connect pairs of annotated spectra. After filtering for structurally unique paired molecules, in total 30 184 annotated spectrum pairs could be extracted from the global molecular network. As biotransformations are reversible, each such pair yields two possible anchor–suspect pairs, with each compound within the pairs in turn serving as an anchor while the other serves as a suspect. Consequently, we were able to extract 60 368 high-quality anchor–suspect pairs.

### Applying relevant biotransformation rules

Once an anchor molecule is identified, BAM applies relevant biotransformation rules. These rules can be sourced from databases like RetroRules [7], KEGG [8], or Metacyc [17]. In this case, RetroRules was used due to its comprehensive collection of biotransformation rules. To extract and apply these rules, we employed PROXIMAL2 [9]. This tool pairs substrate and product molecules based on molecular similarity, aligns each pair using a maximum common substructure algorithm to identify the reaction center, and then creates a lookup table entry defining the changes at the reaction center and its neighbors. These entries, encoded using KEGG atom types [18], are generated for all substrate–product pairs. Finally, for a given anchor–query pair, BAM applies the rules whose mass difference matches the observed precursor mass difference between the anchor and query spectra to generate candidate derivatives for the query molecule.

From the 351 704 biotransformation rules in RetroRules, 84 614 reactant–product pairs mapping to 42 307 unique rules were extracted using PROXIMAL2. When matching rules to observed precursor mass differences, both rule-based and MS/MS mass differences were rounded to unit resolution. Observed precursor mass differences matched 17 254 biotransformation rules, corresponding to 41 % of rules from the RetroRules database that are applicable for BAM. Conversely, 60 320 out of 60 368 anchor–suspect pairs exhibit a mass difference corresponding to at least one biotransformation rule, indicating that almost all queries could potentially be explained by a rule from the RetroRules database. The remaining 48 anchor–suspects pairs that cannot be explained by these rules can be attributed to either an unknown biotransformation or a non-biochemical relationship between the anchor and suspect molecules.

### Ranking candidate structures

The final step involves ranking the generated candidate structures. Multiple candidates can be generated for each anchor–query pair because biotransformation rules might be applied to different atoms in the anchor molecule or because multiple rules match the observed precursor mass difference. Therefore, a systematic approach is needed to prioritize the suggested chemical identities of the suspect. For this, we use GNN-SOM [10], a tool that employs a graph neural network to predict the likelihood of each atom in a molecule being the target of a biotransformation operation. GNN-SOM was trained on enzymatic interactions from the KEGG database to classify each atom within a graph representation of a molecule as the site-of-metabolism for a given enzyme. BAM uses GNN-SOM to rank the generated derivatives based on the likelihood of their corresponding site-of-metabolism, with each derivative being assigned the likelihood of the specific atom to which the biotransformation rule was applied.

The quality of ranking is assessed by determining the rank of the true suspect among all candidates. To resolve ties, the average rank is computed by summing all ranks, including duplicates, and dividing by the number of tied candidates. For example, if three candidates are tied for first place, the average rank would be calculated as 2, and all three candidates would be assigned this rank. Additionally, we report the percentage of true suspect identities ranked at position *k*, where *k* ranges from 1 to the maximum number of unique candidates.

The average number of unique derivatives generated by the previous step is 3.65 (*±*4.29). Among these candidates, GNN-SOM achieves an average rank of 2.1, with 57.7 % of candidates ranked at 1 and 77.2 % of candidates ranked at 2.

### Software availability

BAM was implemented using two different code environments. First, PROXIMAL2 was applied and its results were analyzed using Python 3.9, PubChemPy (version 1.0.4) [19], BioServices (version 1.11.2) [20], scikit-learn (version 1.1.2) [21], NumPy (version 1.23.2) [22], NetworkX (version 2.5) [23], RDKit (version 2022.03.5) [24], KCF-Convoy (version 0.0.5) [25, 26], Pandas (version 1.4.3) [27, 28], Matplotlib (version 3.5.3) [29], and Seaborn (version 0.12.2) [30]. Second, GNN-SOM was applied using Python 3.10, RDKit (version 2022.03.5) [24], PyTorch (version 1.12.1) [31, 32], and PyTorch Geometric (version 2.1.0) [33]. All code, examples files, and detailed instructions are available as open source under the MIT license at https://github.com/HassounLab/BAM.

## Acknowledgments

This research is supported by the NIGMS of the National Institutes of Health, Awards R35GM148219 and R35GM148219. WB was supported in part by the University of Antwerp Research Fund. The content is solely the responsibility of the authors and does not necessarily represent the official views of the NSF nor the NIH.

